# Protective role of the vulture facial and gut microbiomes aid adaptation to scavenging

**DOI:** 10.1101/211490

**Authors:** M. Lisandra Zepeda Mendoza, Gary R. Graves, Michael Roggenbuck, Karla Manzano Vargas, Lars Hestbjerg Hansen, Søren Brunak, M. Thomas P. Gilbert, Thomas Sicheritz-Pontén

**Author notes:** Correspondence to: Thomas Sicheritz-Pontén, M. Thomas P. Gilbert, and M. Lisandra Zepeda Mendoza.

## Abstract

**Background:** Vultures have adapted the remarkable ability to feed on carcasses that may contain microorganisms that would be pathogenic to most other animals. The holobiont concept suggests that the genetic basis of such adaptation may not only lie within their genomes, but additionally in their associated microbes. To explore this, we generated shotgun DNA sequencing datasets of the facial and gut microbiomes from the black and turkey vultures. We characterized *i*) the functional potential and taxonomic diversity of their microbiomes, *ii*) the potential pathogenic challenges they face, and iii) elements in the microbiome that could play a protective role to the vulture’s face and gut.

**Results:** We found elements involved in diseases, such as periodontitis and pneumonia (more abundant in the face), and gas gangrene and food poisoning (more abundant in the gut). Interestingly, we found taxa and functions with potential for playing health beneficial roles, such as antilisterial bacteria in the gut, and genes for the production of antiparasites and antiinsectisides in the face. Based on the identified phages, we suggest that phages aid in the control, and possibly elimination as in phage therapy, of microbes reported as pathogenic to a variety of species. Interestingly, we also identified *Adineta vaga* in the gut, an invertebrate that feeds on dead bacteria and protozoans, suggesting a defensive predatory mechanism. Finally, we suggest a colonization resistance role though biofilm formation played by *Fusobacteria* and *Clostridia* in the gut.

**Conclusions:** Our results highlight the importance of complementing genomic analyses with metagenomics in order to obtain a clearer understanding of the host-microbial alliance and show the importance of microbiome-mediated health protection for adaptation to extreme diets, such as scavenging.

## Background

Vultures are scavengers whose global populations are under serious threats, and a better understanding of various aspects for their biology are necessary [1,2]. Vultures are known as nature’s clean-up crew as they feed on muscles and viscera from carcasses of animals that have died mainly from malnutrition, accidents, and natural or infectious diseases, thus are expected to carry pathogens such as those causing anthrax, tuberculosis, brucellosis, etc. [3]. Carcasses are a very nutrient-rich resource and it has been speculated that the release of toxins and pathogenicity genes in the carcass microbiome are part of a microbial strategy for outcompeting other microbes [4,5]. The main colonizers of a carcass are microbes originating from the normal flora of the dead animal, which might turn pathogenic in the carcass environment [6], as well as soil-dwelling bacteria, nematodes, fungi and insects [7]. In spite of the potentially serious health implications posed by their consumption, the pathogenic repertoire of the gut of these species of birds has not been fully characterized in relation to their possible implications in the environment. Thus, one of the most intriguing aspects of vulture biology is how they protect themselves against the health challenges posed by their dietary source. Physiologic, genetic and genomic analyses of different species of vultures have explored this aspect and identified genes associated to respiration, immunity and gastric secretion as candidates for the adaptation to its scavenging diet [8,9].

With the genomic revolution it has become apparent that besides genomic changes, host-associated microbiota play an important role in diet specialization across vertebrates [10] and that the gut microbiome may play highly relevant yet unexplored role in diet-driven speciation [11]. The gut microbiome is related to digestion traits, such as energy harvest, nutrient acquisition, and intestinal homeostasis [12], among other phenotypes related to the immune and neuroendocrine systems [13–15]. Furthermore, the role of the microbiome in health protection to the host has also been demonstrated [16,17], and disorders in it lead to diseases such as irritable bowel syndrome, inflammatory bowel disease, obesity and diabetes [18–20]. In light of the key roles that host-microbiome relationships can play in adaptation, it has been acknowledged that genomic adaptations alone may not provide the full answer to the vulture adaptations for scavenging [8]. However, neither the complete microbial taxonomic diversity (including non-bacterial microbes) nor the gene catalogue of any vulture species’ face and gut microbiomes have been examined for their protective role against microbes that would normally pose serious health risks for other non-scavenging vertebrate species.

In order to evaluate the protective role of the vulture’s facial and gut microbiome, we generated metagenomic datasets from facial swabs and gut samples for two species of New World vultures, the black vulture (*Coragyps atratus*) and the turkey vulture (*Cathartes aura*), and performed metagenomic taxonomic and functional analyses. We identified a wide variety of taxa and genes that would cause serious diseases, e.g. sepsis, gangrene and food poisoning, to animals without protection provided by a specialized scavenging microbiome. Such protection could consist of health defence elements not only from bacterial probiotics, but also from phages and predatory eukaryotes. Our results suggest that the vulture’s microbiota plays a significant health-protective role in its adaptation to a scavenging diet.

## Methods

### Sampling method and DNA sequencing

We analysed a subset of the sample set used by Roggenbuck *et al.* [21]. Briefly, the vultures were collected over a period of several days in Tenessee, USA. *Coragyps atratus* were live-trapped at deer carcasses and then transported to a central facility within a couple of hours of trapping. They were then euthanized with CO_2_, necropsied, and sampled within 30-45 minutes of death. *Cathartes aura* vultures were shot at roosts, bagged individually, and transported to the processing facility where they were refrigerated 2-6 hours before necropsy and sampling. The samples were kept in RNAlater and DNA was extracted and the shotgun libraries for HiSeq PE 100 prepared using the Nextera library building kit following the manufacturer’s instructions, as in Roggenbuck *et al.* [21]. We used a total of 22 turkey vulture (*Cathartes aura*) and 25 black vulture (*Coragyps atratus*) large intestine samples, and 16 turkey vulture and 17 black vulture facial samples.

### Data processing

Two pipelines were used for the processing of the raw reads. In the first approach, we removed adapter sequences and bases with quality <15 using Trimmomatic v0.32 [22]. Afterwards, in order to filter out non-bacterial reads of avian, human, and *Phi* phage origin, the datasets were mapped against the bird genomes dataset of the avian phylogenomic project [23] (which includes the turkey vulture genome), the human (hg19) and the *Phi* phage genomes, and only the non-mapping reads were kept. The second approach was developed in order to take into account the possible *k*-mer bias in the first bases of the reads that could have implications in the subsequent *de novo* assembly and gene prediction. To this end, we trimmed the first 16 bases of the reads with Trimmomatic v0.32 in order to remove *k*-mer bias. Then, we processed those reads with MOCAT [24] to clean them of low quality bases and adaptors and screen them versus *Phi* phage, human and turkey vulture genomes.

### Taxonomic profiling

We used MGmapper [25] to map with bwa v0.7.10 [26] the filtered cleaned reads against the next databases in full mode: MetaHitAssembly [27], HumanMicrobiome [28], ResFinder [29], Plasmid, Virulence, GreenGenes [30], and Silva [31]. We also mapped in chain mode to the next whole genome databases downloaded from GenBank in the given order: human, plants, vertebrates, invertebrates, protozoa, fungi, and viruses. The remaining non-mapping reads were mapped in chain mode to the whole genome databases of bacteria. The MGmapper results were used to obtain rarefaction curves from each dataset using an in-house script. Using the unique mapping reads, we calculated the coverage (percentage of reference sequence covered by reads) of each identified species and filtered based on it as follows:

1. We used a relaxed filtering in order to ensure the identification of low abundant taxa, in which we removed identifications with 90% of the abundance signal coming from only 3 samples.
2. We used a stricter filtering in which, on top of removing those with a low signal of abundance, we kept the species identified with more than the 1^st^ quartile (Qu) coverage from the coverage distribution of the corresponding database. With the filtered taxa, we identified those species present in at least 90% and 50% of all the samples, thus defining a relaxed and a stricter taxonomic shared face and gut microbiome core.

For each database, we compared the taxa present only in the facial and gut datasets, those present in both, and those in significantly different abundance (*P*< 0.05). To identify the differentially abundant species, we performed a t test on the normalized abundance distribution of the species in all the facial dataset versus all the gut dataset. We also evaluated the taxonomic intra and inter samples variation between the facial and gut samples by calculating the Euclidean distances of their normalized abundances using the ward.D method in R [32].

As a second approach, we used only the unique mapping reads from MGmapper and kept the taxonomic identifications until the species level removing the low abundant ones (90% of the signal coming from less than 3 samples) and normalizing the counts. We used these assignations to test for microbial abundance correlation by calculating the Spearman correlation for each pairwise comparison of the microbes and calculated the p values (*P*) with a Bonferroni correction on those with a correlation value > 0.8 and < -0.7. We also examined the enrichment and depletion of taxa within the face and gut microbiome, by getting their mean abundance across the samples and comparing to the total distribution to calculate the Bonferroni corrected *P*. From these assignations, we also obtained a specific face and gut core. We defined two types of cores: a strict one in which we keep those present in at least 80% of the samples, and a relaxed one in which we keep those present in at least 50% of the samples.

We also identified the taxa of the top most abundant proteins (those with more than 2,000 mapping reads in the face dataset and 5,000 in the gut dataset) and analysed their principal components (PCs) and their rotation matrix to identify which taxa drive the face and gut microbiota intra-samples variation. We defined as “variation drivers” those with an absolute rotation matrix value larger than the 3^rd^ Qu value of the distributions from PC1, PC2 and PC3, and as “non-variation drivers” those with less than that threshold.

Besides the MGmapper identifications, we used MOCAT as a third taxonomic identification method. For this approach we used the MOCAT strict non-redundant gene catalogue taxonomic annotation, with minimum length of 80 amino acids, no low abundant, only bacteria, fungi and virus and with a Uniprot annotation (see Methods - Functional profiling), which we analysed with MEGAN [33] using as input the search against Uniprot using usearch [34]. We used MEGAN to annotate the microbial attributes and to compare their normalized abundance in the gut and face microbiomes.

### Pathogenic characterization

We looked for potential pathogens in the filtered bacterial and plasmid identifications. To this end, we downloaded a list of the bacteria annotated with a disease from the database Pathosystems Resource Integration Center (PATRIC) [35]. In PATRIC, bacteria are annotated as pathogenic if they have been reported with experimental data as causative of a disease in a species. We further added the pathogenicity classification level of bacterial strain using the list from van Belkum [36], which was developed by the Commissie Genetische Modificatie (COGEM). The pathogenicity classes are defined as follows. Class 1 represents species that are commonly non-pathogenic, although there may be differences in virulence between the species strains that should be taken into account. Class 2 contains species that can cause diseases in humans or animals and that are unlikely to spread in the human population. And class 3 encompasses species that cause serious human diseases and can disseminate in the human population. By using the metadata of the pathogenic strains downloaded from PATRIC, we also identified whether or not the bacteria are capable of sporulation and of antimicrobial resistance, together with the disease and the reported host. For the identification of pathogenic plasmids, we used the list reported by Ho-Sui *et al.* [37], where they analysed the association of virulence factors with genomic islands from pathogenic bacteria.

We used R v3.1.1[32] to examine the distribution of the total number of identified pathogenic bacteria. We grouped the samples by *i*) vulture species (turkey and black vulture), and *ii*) body sampling place (face and gut). We then tested with a two-tailed and one-tailed (alternate greater) t test if the means of the distributions were significantly different. Next, we examined the number of samples in which each pathogenic bacterial strain, plasmid, resistance gene, and virulence factor was present. In order to get a potentially pathogenic core, we identified those in 50% and 90% of the samples and compared those unique to the face and the gut.

### Abundance analyses of pathogenic bacteria

In order to analyse the abundance of the potentially pathogenic microbes across the samples, we first rescaled the number of unique mapping reads to bacteria by their percentage in the sample. Afterwards, bacteria present in low abundance (those with 90% of their signal coming from 3 or less samples) were removed. In order to see if the samples differentiate by sampled body part (face or gut), we used the rescaled counts to build a dendrogram using a hierarchical clustering on the Euclidean distance. Afterwards, we examined which of the retained potentially pathogenic bacteria were present only in the face or in the gut, and which ones were in both. We then used a t test to evaluate if the abundance of those present in both face and gut was statistically different by sampling body place and by vulture species.

### 16S bacterial taxonomic comparison

We compared the taxonomic bacterial identifications from both gut and facial datasets obtained by 16S analyses by Roggenbuck *et al.* [21] against the bacterial identifications from our metagenomics datasets using MGmapper and the taxonomic annotation of the *de novo* assembled genes with Uniprot. Afterwards, we also used the taxonomic identifications obtained from the unmapped reads with DIAMOND [38] for comparison.

### Functional profiling

Using the reads mapping to the databases of interest with MGmapper and the unmapped reads we performed *de novo* assembly with IDBA-UD [39] and predicted genes with Prodigal [40]. Afterwards, we generated a non-redundant (nr) gene catalogue with usearch [34] by clustering the predicted genes with 90% identity and keeping the centroid sequences. Next, the nr gene catalogue was searched with ublast [34] against Uniprot [41], and the resulting identifications were functionally and taxonomically annotated with the use of a customized python script. Finally, we used DIAMOND v0.6.4 [38] blastx to search the unmapped reads against Uniprot, keeping only the best hit for subsequent functional and taxonomic annotation. We kept as part of the functional core those genes present in more than a given number of samples according to their presence distribution.

For further assessment at a functional level, we converted the Uniprot ids to the KEGG [42] E.C id and its corresponding pathway. Using the classified proteins, we built a matrix for performing principal component analyses (PCA). Looking at the rotation matrix from the PCA we identified those pathways with an absolute rotation value of the 1^st^, 2^nd^, and 3^rd^ PC within the minimum and 1^st^ Qu values of their distributions. In order to identify which pathways drive most of the variation between the face and gut microbiomes, we identified those pathways for which the absolute rotation value of their 1^st^, 2^nd^, and 3^rd^ PCs was larger or equal to the 3^rd^ Qu value of their distributions. To test whether the functional profiles of the face and gut microbiomes are similar, in spite of having large intra sample type variation, we performed replicates of down sampling the number of proteins corresponding to each class pathway by the minimum abundance of the distribution. We also obtained the Euclidean distances on the rescaled values of the matrices used for the PCAs.

As a second method, we used MOCAT with SOAPdenovo v1.05 [43] for *de novo* assembly with the reads cleaned with the second approach. Subsequently, we corrected the assembly for indels and chimeric regions with SOAPdenovo. Using prodigal we then predicted the genes from all the samples, pooled them, and built an nr gene catalogue with uclust [34] using a 90% identity threshold. This was done separately for the face and gut datasets. In the MGmapper core definition approach, the nr gene catalogue was obtained for each sample, then the catalogues were pooled and the unique genes were kept to compare their presence or absence across the samples. In contrast, in this approach using MOCAT, we built the cores based on the abundance of the reads mapping to the nr gene catalogue. To this end, we first mapped the reads of each sample against the nr gene catalogue, then rescaled the counts values, removed the low abundant genes (genes with less than 200 reads mapped), those without a Uniprot annotation, and those not derived from bacteria, archaea, virus, or fungi. Furthermore, the proteins must have had at least 80 amino acids aligned to the Uniprot hit for its annotation. From these proteins, we also obtained a strict (at least 80% of the samples) and a more relax (at least 50% of the samples) core. We also identified the top most abundant proteins (those with more than 2,000 mapping reads in the face and 5,000 in the gut samples). On the relaxed functional core, we performed pathway functional analyses of their E.C. ids with KEGG.

### Antibiotic resistance

In order to search for antibiotic resistance genes, besides searching the ResFinder database with MGmapper, we downloaded the Resfams v1.2 database [44], a curated database for antibiotic resistance genes, and the associated profile hidden Markov models. We searched the *de novo* assembled nr gene set of each sample against the Resfams profiles using hmmscan of HMMER v3.0 [45].

### Damage pattern

To test for DNA damage caused by the gastrointestinal acidic conditions of the vulture, we used the reads mapped against the nr gene sets from the face and gut datasets and used them as input for MapDamage [46], which calculates the nucleotide misincorporations in the 5’ and 3’ extremes of the DNA reads.

## Results

### Metagenomic dataset

From the total of 48 different individuals (25 *Coragyps atratus*, and 23 *Cathartes aura*), we used 33 facial samples (17 *Coragyps atratus*; 16 *Cathartes aura*) and 47 intestinal samples (*25 C. atratus*; 22 *C. aura*). We produced a total of 342,279,763 raw read pairs from the facial samples and 512,803,778 from the gut samples. After cleaning and removing endogenous DNA by mapping against the bird genomes of the avian phylogenomic project [23], we obtained 79,938,910 read pairs from the facial samples (with a median of 1,378,000 read pairs per sample) and 144,877,366 from the gut samples (with a median of 1,118,000 per sample) (Additional File 1, Table S1).

To prove the consistency of the taxonomic profiling between the two species, we compared the number of identified taxa in each species. We filtered the MGmapper [25] identifications of each whole-genome database by depth and breadth (percentage of covered reference sequence) of coverage and identified taxa differentially abundant in the face and gut samples (Tables S2, S3). The number of identified bacteria is not significantly different between vulture species (*P*= 0.52, *C. atratus* mean= 366.97, *C. aura* mean= 334.65). Neither for fungi (*P*= 0.43, *C. atratus* mean= 9, *C. aura* mean= 7.5), viruses (*P*= 0.33, *C. atratus* mean= 21, *C. aura* mean= 28.2), plasmids (*P*= 0.68, *C. atratus* mean= 186.85, *C. aura* mean= 173.65), and protozoa (*P*= 0.21, *C. atratus* mean= 12, *C. aura* mean= 9.62). Also, the number of identified proteins with resistance to antibiotics is not different between vulture species (*P*= 0.64, *C. atratus* mean= 107.5, *C. aura* mean= 100.78). We compared our metagenomic bacterial identifications to those by Roggenbuck *et al.* [21]. A total of 735 bacteria were identified by 16S, out of which 93 are not found among our metagenomics identifications with strict filtering. When using the pre-filtering identifications and those identifications from the Silva and GreenGenes, only 14 genera were not identified (Additional File 1).

### Taxonomic characterization

Compared to the gut, the facial microbiome has higher microbial diversity in terms of number of taxa and variation between individuals (*P* protozoa= 0.021, *P* fungi= 0.029, *P* bacteria= 0.000196) (Figures S1-S3), however, there is no significant difference in abundance (*P* protozoa= 0.514, *P* fungi= 0.47, *P* bacteria= 0.71). Although the number of identified virus is not significantly different between face and gut (*P*= 0.58), they are statistically less abundant and variable in the face than in the gut microbiome (*P* abundance= 0.00017, face Euclidean distance= 9.04, gut Euclidean distance= 13.14). Most of the bacteria present in higher abundance in the face microbiome belong to *Pseudomonas*, *Bacteroides*, and *Prevotella*, while most of those present in higher abundance in the gut microbiome are *Escherichia*, *Campylobacter*, and *Clostridium* (Additional File 2, Figure S4).

We identified a total of 143 bacterial strains significantly more abundant in the face microbiome, 46 after the breadth filtering (mostly belonging to *Pseudomonas*). In the gut, we identified 56 bacterial strains, 33 after the breadth filtering (mostly belonging to *Escherichia* and *Campylobacter*). Those more abundant in the face dataset can be classified into *i*) potential pathogens, *ii*) related to bioremediation (ionizing resistant, reducers of heavy metals, or oil degraders), and *iii*) potentially beneficial (producers of antibiotics, insecticides and antifungals), usually intestinal bacteria, and related to water, plants, or soil. And those significantly more abundant in the gut dataset can be classified as *i*) potential pathogens, *ii*) potentially beneficial, usually intestinal or faecal bacteria from chicken, and *iii*) fermenters and producers of intestinal metabolites or intermediary metabolic substances.

From the mapping of the reads to the nr gene set catalogues obtained from MOCAT, the taxa of the top most abundant proteins in the face microbiome are from *Sporidiobolales* (fungi), *Orthoretrovirinae* (virus), *Pleosporaceae* (fungi), *Bacillus cereus*, *Streptococcus*, and *Clostridiales*. While in the gut microbiome they are from *Bordetella*, *Mycobacterium*, *Chlamydia*, *Clostridium*, *Blautia* (a species identified in the mammalian gut [47,48]), and *Carnobacterium* (certain species have preservative qualities of meat products by inhibiting the growth of *Listeria monocytogenes* [49,50]). The results from the search against Uniprot analysed with MEGAN (Figs. 1, 2) show that in the face microbiome the dominant population is *Proteobacteria*, followed by *Bacteriodetes*, *Firmicutes*, and *Actinobacteria*, with *Fusobacteria* in 12^th^ place. Deeper examination of the *Proteobacteria* from the facial microbiome shows that the most abundant taxa are *Burkholderiales* from the *Betaproteobacteria*, and *Pseudomonadales* from the *Gammaproteobacteria* (Fig. 1B). Inside the *Pseudomonadales*, the most abundant taxon is *Psychrobacter* (mainly *P. cryohalolentis*, and *C. articus*), followed by *Pseudomonas* (mainly *P. stutzeri*, *P. aeruginosa*, and *P. putida*) (Fig. 1D-F). From the *Bacteroidetes*, the most abundant taxa are *Prevotellaceae* (mainly *P. ruminicola*) and *Flavobacteriaceae* (mainly from unclassified *Flavobacteriaceae*, followed by *Flavobacterium* genera) (Fig. 1C).

**Fig. 1.**
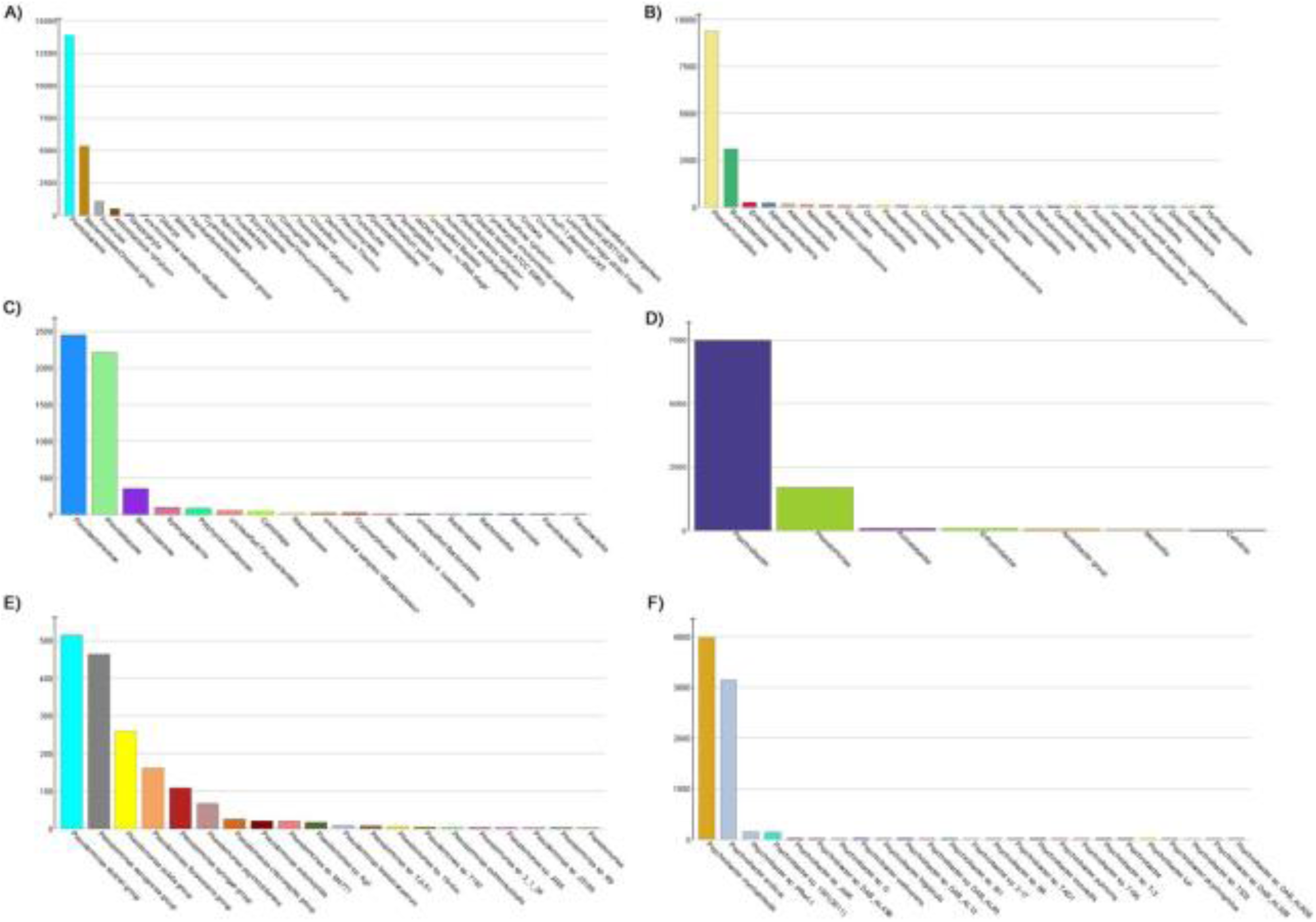
Taxonomic profile of the facial microbiota with MEGAN filtered nr gene catalogue. (**A**) Phylum level, (**B**) Proteobacteria, (**C**) Bacteroidetes, (**D**) Pseudomonadales, (**E**) *Pseudomonas*, and (**F**) *Psychrobacter*.

**Fig. 2.**
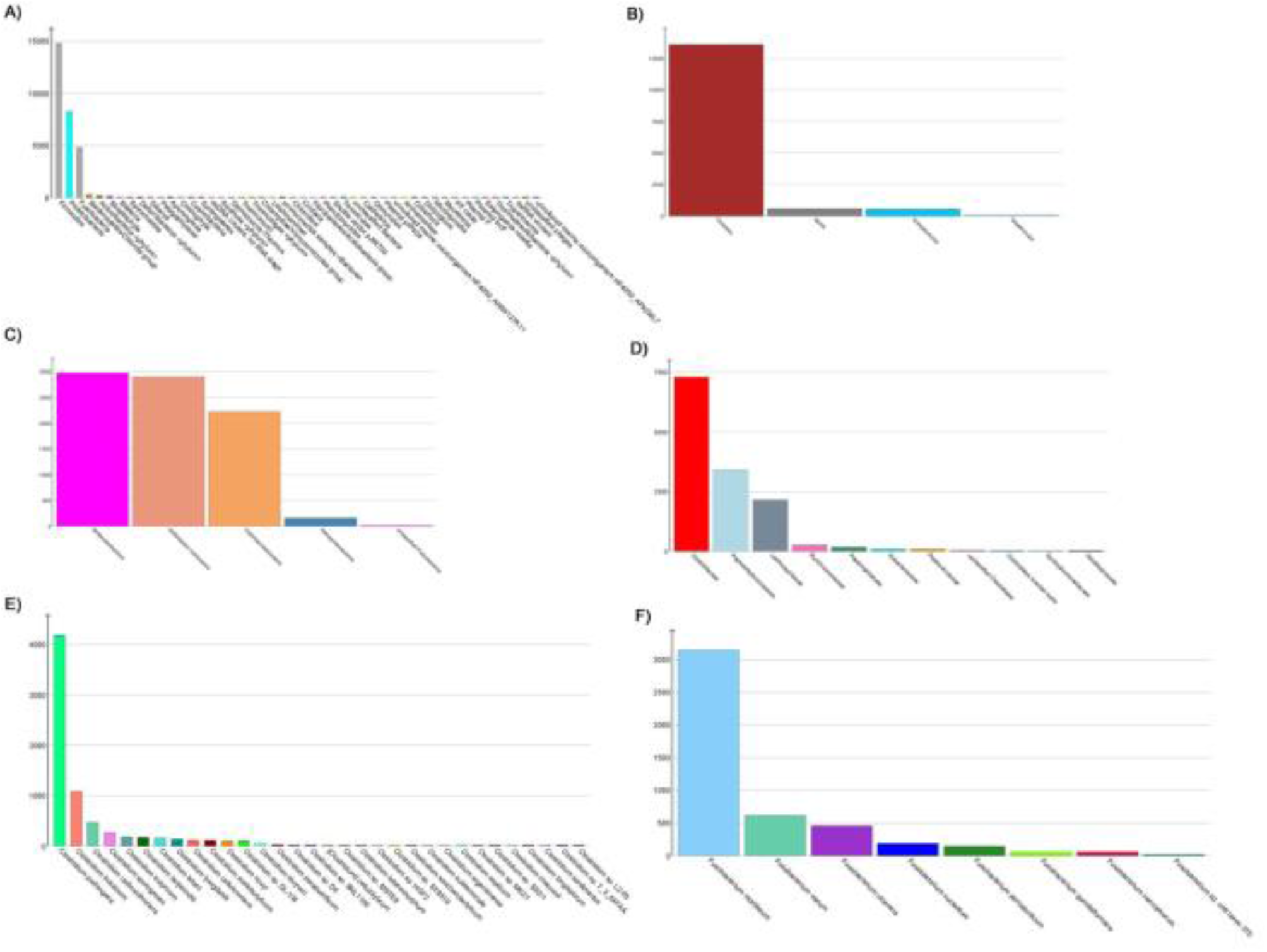
Taxonomic profile of the gut microbiota with MEGAN filtered nr gene catalogue. (**A**) Phylum level, (**B**) Firmicutes, (**C**) Proteobacteria, (**D**) Clostridiales, (**E**) *Clostridium*, and (**F**) *Fusobacterium*.

The gut microbiome consists of *Firmicutes* as the most abundant phyla, followed by *Proteobacteria*, *Fusobacteria* in the third place, and much less abundant *Bacteroidetes* in the fourth place. Deeper examination of the gut microbiome *Firmicutes* shows that the most abundant taxa are *Clostridia* (Fig. 2B), among the *Clostridiales*, the most abundant taxon is *Clostridiaceae*, followed by *Peptostreptococcaceae* and *Lachnospiraceae* (Fig. 2D). From the *Clostridium*, the most abundant taxa are the potential pathogens *C. perfringes* and *C. butilinium*, followed by the beneficial *C. carboxinovorans*, *C. sporogenes*, and *C. butyricum* (Fig. 2E). From the *Proteobacteria*, the most abundant taxa are *Burkholderiales* from the *Betaproteobacteria*, *Epsilonproteobacteria* from the delta/epsilon subdivision, and *Enterobacteriales* (mainly from *Escherichia*) from the *Gammaproteobacteria* (Fig. 2C). Looking at the *Fusobacteria*, the most abundant taxa are the potential pathogens *Fusobacterium mortiferum*, *F. varium*, and *F. ulcerans*, followed by oral related *Fusobacterium* (Fig. 2F).

Many *Clostridia*, usually part of a normal human gut microbiome, are found in the vulture gut, likely playing relevant digestive roles (Additional File 3). For instance, we identified as significantly more abundant in the gut *C. saccharolyticum*, a sewage sludge isolated bacteria that ferments various carbohydrates into acetic acid, hydrogen, carbon dioxide, and ethanol [51], functions for which we identified related genes. Also, the cellulose degraders *C. cellulovorans* and *C. lentocellum* [52] are significantly more abundant in the gut microbiome, for which we identified genes related to cellulose degradation. The vulture’s gut microbiome also contains *Bacteroides xylanisolvens*, which breaks down xylan in the human gut [53] and for which we identified a gene related to this function. Also, significantly more abundant in the gut microbiome than in the face are *C. beijerinckii*, which produces butanol, acetone and isopropanol using various substrates such as pentoses, hexoses and starch [54], and *C. saccharobutylicum*, a butanol and ethanol-producing bacteria [55].

We also identified protein coding genes involved in vitamin biosynthesis in the gut strict MOCAT nr functional core. For example, we identified D-threo-aldose 1-dehydrogenase, involved in ascorbate and aldarate metabolism, cobalamin biosynthesis, riboflavin biosynthesis, thiamine biosynthesis (vitamin B_1_), 2-ketopantoate reductase, involved in vitamin B_5_ production [56], from the genera *Hydrogenophaga*, *Herbaspirillum*, and *Gordonia*. In the MGmapper second type of core, that of gene core not taking the taxa into account, we identified a larger abundance of genes annotated to belong to the metabolism of cofactors and vitamins than in the face, such as folate biosynthesis, vitamin B_6_ metabolism, riboflavin metabolism, and retinol metabolism. We also identified genes for the biosynthesis of various essential amino acids.

### Comparison of the facial and gut microbiome variation

Based on the PCA on the abundance of the identified species in the face and gut datasets (Fig. 3AB), we found that 117 of the 803 species identified as driving the variation in the face microbiome are pathogenic bacteria to a mammalian host (e.g. species from the genera *Bordetella*, *Gordonia*, *Shigella*, *Yersinia*, *Brucella*, *Prevotella*, and *Treponema*), soil or plant related, and some related to mucosal surfaces and normal oral flora (*Neisseria* and *Nocardia*). The 406 species not driving most of the variation include genera related to bioremediation (e.g. *Acinetobacter*), as well as other pathogens (*Arcobacter* and *Brucella*). Phages of *Clostridium*, *Pseudomonas*, *Shigella*, and *Staphylococcus* are among those non-variant, and phages of *Samonella*, *Aeromonas*, and *Erwinia* are among the variant ones.

**Fig. 3.**
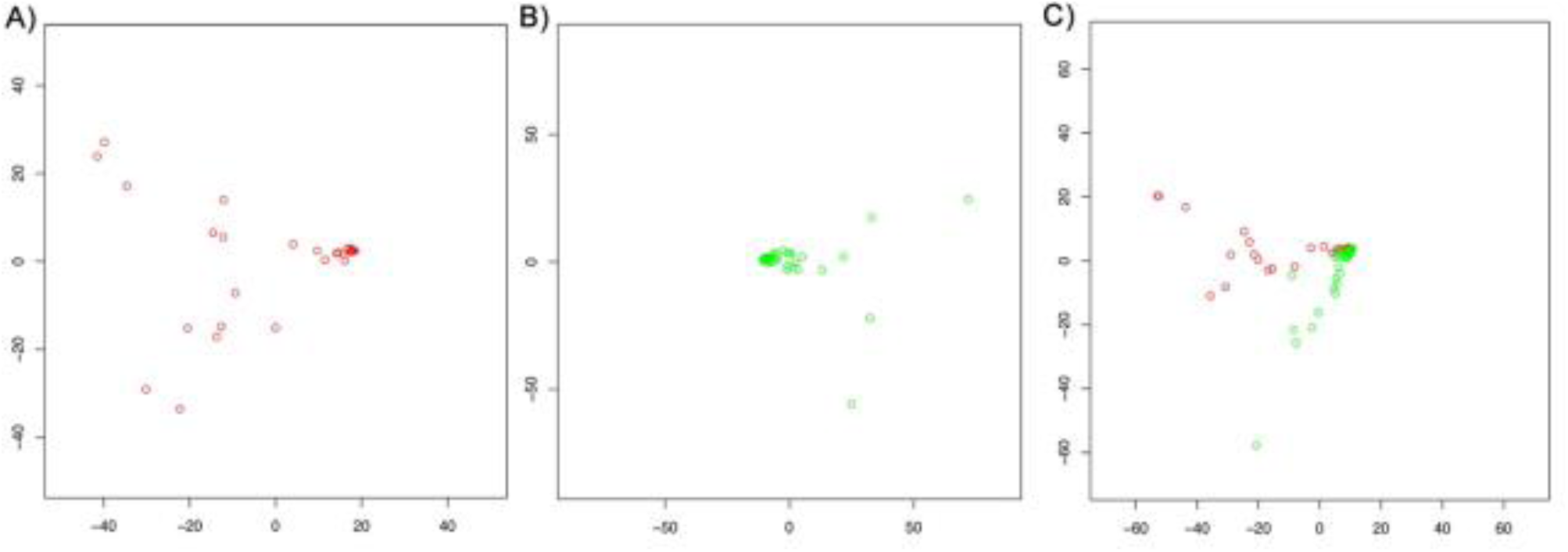
PC1 vs PC2 of the taxonomic species level abundance. (**A**) Face microbiome, (**B**) gut microbiome, and (**C**) species present in both face (red) and gut (green).

In the gut samples, we identified 604 species that were significantly variant between the samples and 348 as not variation drivers. The variation drivers include 112 potentially pathogenic bacteria, such as species from the genera *Listeria*, *Shigella*, *Yersinia*, *Bordetella*, *Shewanella*, *Erwinia*, and *Vibrio*. And the non-drivers include species from the genera *Escherichia*, *Bacillus*, *Brucella*, and *Clostridium*, among others. Non-variation driver phages include phages for *Escherichia*, *Enterobacteria*, and *Shigella*. While those driving variation include phages for *Clostridium*, *Yersinia*, and *Pseudomonas*.

Examining the 879 microbial species shared by the face and gut samples (Fig. 3C), we identified 553 species as driving variation (62.9%), and 326 as non-variation drivers (37.1%). Among the variation drivers are *Yersinia*, *Ralstonia*, *Rhizobium*, *Bifidobacterium*, *Bordetella*, *Listeria*, and *Burkholderia*. Those non-variation drivers include *Brucella*, *Treponema*, *Clostridium*, and *Campylobacter*. Looking at the phages, only *Pseudomonas* phages are variation drivers, while non-variation drivers include phages for *Clostridium*, *Enterobacteria*, *Erwinia*, and *Shigella*.

### Functional potential characterization

PCA of the MGmapper nr gene set catalogue of the pooled face and gut microbiomes annotated with KEGG shows less variation than the taxonomic profile. The PC1 of the analysis of each pathway class explains 78%-99% of the variance. From the PC1vsPC2 plots, it is clear that PC1 separates the samples by their functional composition, which is similar, while PC2 separates them by their sampling origin (face or gut) (Fig. 4, Figure S5). To test whether the profile of the functional potential of the face and gut microbiomes are very similar in spite of the large intra sample type variation, we down sampled the number of proteins corresponding to the given class pathway by the minimum of the distribution. The resulting down sampled PCAs show a reduced scale in the PC1 while the PC2 scale does not change and the samples largely overlap (Fig. 5), confirming our observation of functional similarity in spite of variation.

**Fig. 4.**
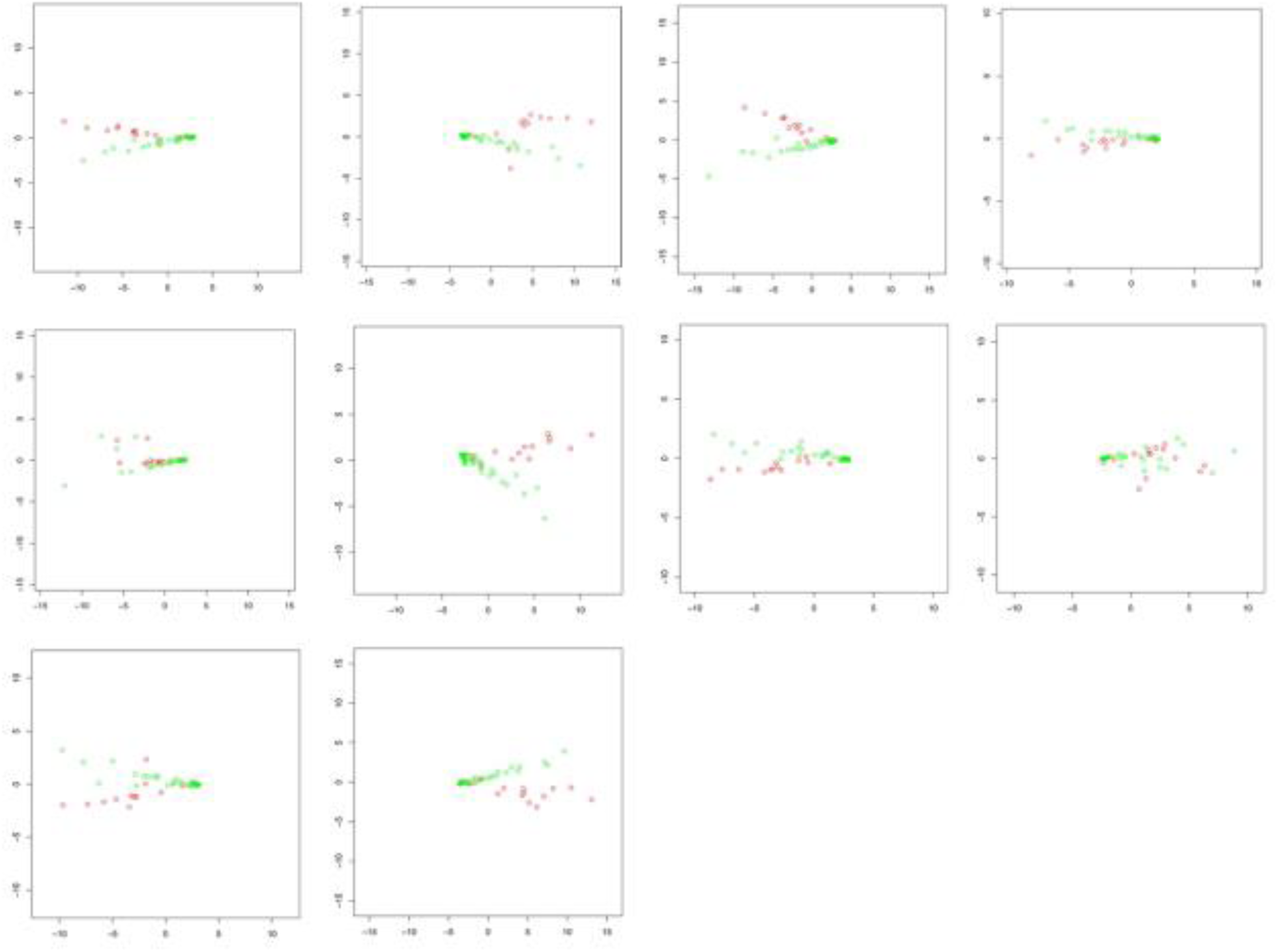
PC1 vs PC2 of each pathway class. Red dots are samples from the face dataset, green dots are from the gut dataset. The pathway classes examined are (left to right, top down): amino acid metabolism, metabolism of other secondary metabolites, carbohydrate metabolism, energy metabolism, glycan biosynthesis and metabolism, lipid metabolism, metabolism of co-factors and vitamins, metabolism of other amino acids, metabolism of terpenoids and polyketides, and xenobiotics biodegradation and metabolism.

**Fig. 5.**
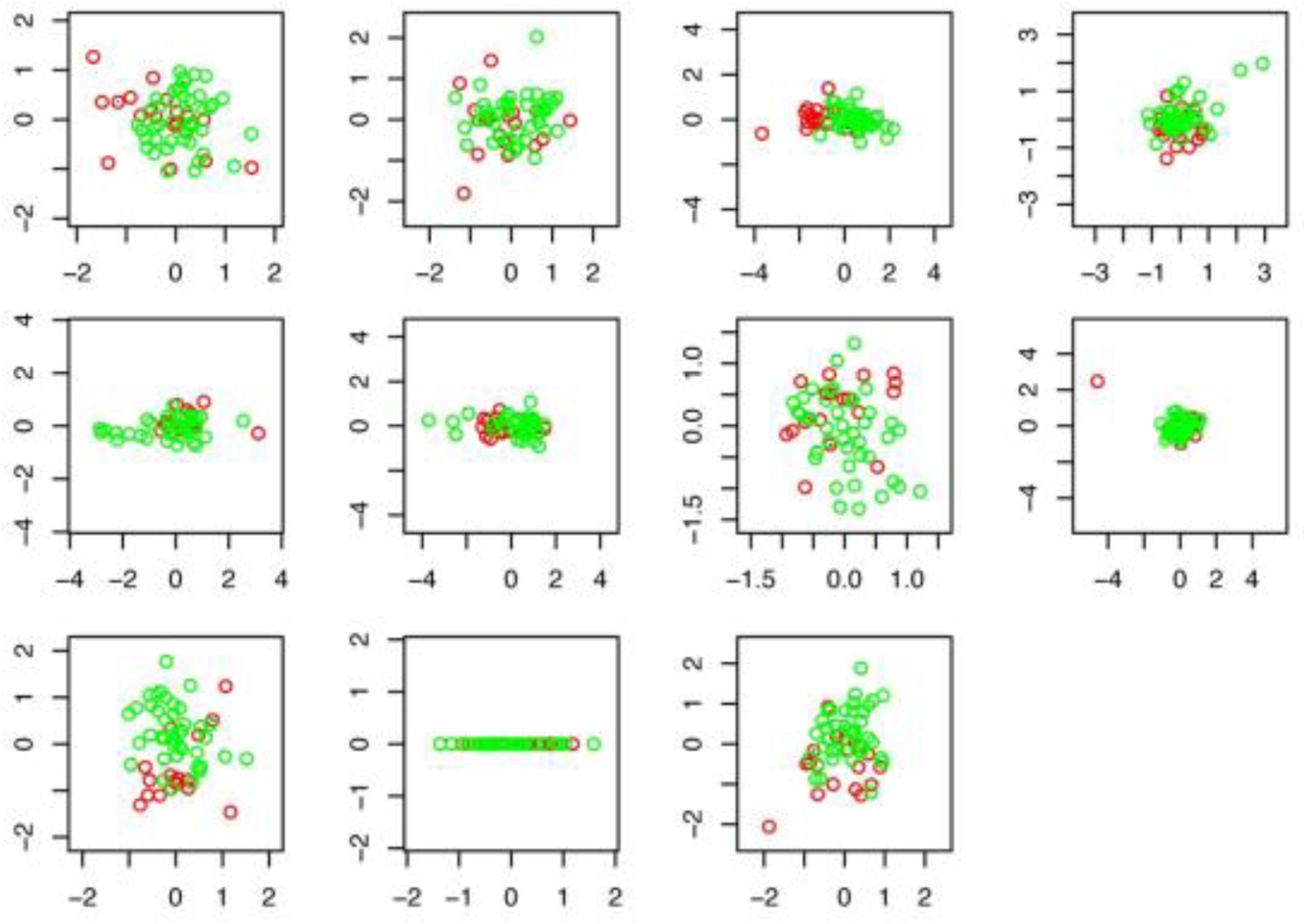
PC1 vs PC2 of each pathway class with down sampled protein counts. Red dots are from the face samples, green dots are from the gut datasets. The pathway classes examined are (left to right, top down): Amino acid metabolism, metabolism of other secondary metabolites, carbohydrate metabolism, energy metabolism, glycan biosynthesis and metabolism, lipid metabolism, metabolism of co-factors and vitamins, metabolism of other amino acids, metabolism of terpenoids and polyketides, and xenobiotics biodegradation and metabolism.

Looking at the rotation matrix from the PCA, 60.4% (87) of the sub-pathways drive variation between the face and gut microbiomes. Analysing the sample type intra variation, 57% of the sub-pathways in the face (81) and 55.94% (80) in the gut datasets drive intra sample type comparison. We identified 59 genes in the top 5% variation driver genes with the largest mean difference in abundance between face and gut. Of those genes, 18 correspond to amino acid metabolism (17 only present in gut samples and 1 only in one face sample), 15 genes from carbohydrate metabolism (all of them only in gut samples), and 7 genes corresponding to metabolism of cofactors and vitamins (all only in gut samples). We found that 44% of the genes (921 out of the KEGG annotated 2093 pooled face and gut relaxed gene cores) do not drive variation. The 46 genes of the top 5% with the smallest mean difference in abundance between face and gut correspond to the metabolism of amino acids, carbohydrates, cofactors and vitamins, glycan biosynthesis and metabolism, lipid metabolism, energy metabolism, chlorocyclohexane and chlorobenzene degradation, metabolism of terpenoids and polyketides, and metabolism of other amino acids.

Based on the Euclidean distance measures, the gut has less intra sample type variation than the face (Table 1, *P*= 0.002). The face microbiome distances range from 3.2 to 6.21, while those in the gut microbiome range from 2.175 to 3.676. Analysis of all the functions together (Figure S6) instead of per pathway class (Fig. 5, Figure S7), shows that the face and gut microbiomes clearly separate into 2 different clusters (PC1= 0.847, PC2= 0.075). The MOCAT functional characterization yielded a total of 38,403 nr genes from the face dataset, and 50,106 nr genes from the gut dataset. Based on the normalized abundance of the mapping reads, we identified 1,507 genes in the facial strict core and 7,215 in the relaxed, and 157 very abundant ones (with more than a total of 2,000 mapping reads). We found 2,512 genes in the gut strict core, 14,028 in the relaxed one, and 151 top abundant ones (with more than 5,000 mapping reads).

**Table 1.** Table 1. Distances between and within the face and gut samples.

| Pathway | FaceVsFace | GutVsFace | GutVsGut |
| --- | --- | --- | --- |
| Amino acid metabolism | 4.868 | 4.663 | 2.865 |
| Biosynthesis of other secondary metabolites | 5.97 | 5.972 | 3.676 |
| Carbohydrate metabolism | 4.099 | 4.479 | 3.581 |
| Energy metabolism | 3.334 | 3.269 | 2.175 |
| Glycan biosynthesis and metabolism | 3.240 | 3.675 | 3.22 |
| Lipid metabolism | 5.601 | 5.662 | 3.035 |
| Metabolism of cofactors and vitamins | 4.273 | 4.410 | 2.992 |
| Metabolism of other amino acids | 3.484 | 3.767 | 2.753 |
| Metabolism of terpenoids and polyketides | 4.566 | 4.936 | 3.184 |
| Nucleotide metabolism | 1.497 | 1.559 | 1.342 |
| Xenobiotic biodegradation and metabolism | 6.217 | 5.981 | 3.613 |

### Core microbiome identification and attributes comparison

In the filtered MGmapper taxonomic profiling, we identified 1,483 species in the facial samples, 638 of which are in at least 50% of the samples (relaxed core), and only 184 in at least 80% of the samples (strict core). In the gut microbiome, we found 1,419 microbial species, with only 322 are present in at least 50% of the samples, and 129 in at least 80%. In the functional characterization, we identified a total of 238,065 nr unique bacterial genes in the face microbiome and 387,951 nr unique bacterial genes in the gut microbiome (Additional File 4, Tables S4, S5).

We compared the facial and gut taxonomic and functional composition, and the specific microbial attributes from the taxa identified from the annotations of the assembled genes (Additional File 2). We found ~26x more habitat-specialized microbes in the face than in the gut microbiome (face= 8,373, gut= 320), and consistent with the gut anaerobic environment, the gut microbiome has ~5x more anaerobic or microaerophilic bacteria than the face microbiome (gut= 34,749, face= 6,699). The only two functional pathways clearly separated by sample type are lipid and carbohydrate metabolism, followed, to a lesser extent, by biosynthesis of other secondary metabolites, and glycan biosynthesis and metabolism (Fig. 4). From the energy metabolism class, the methane metabolism is among the most abundant functions in the face and gut microbiomes (Figure S7).

### Pathogenic characterization

There is no significant difference in the number of identified potentially pathogenic plasmids between the face and gut microbiomes (*P*= 0.44, face mean= 11.3, gut mean= 9.82) (Figure S8A), and there is no statistical difference in their abundance (*P*= 0.775). Furthermore, there is no potentially pathogenic plasmid present in 90% of the samples. Among those present in at least 50% of the samples are plasmids from *Burkholderia vietnamiensis*, *Escherichia coli*, and *Ochrobactrum anthropic*. The Shiga toxin 1-converting phage BP-4795, which transmits virulence genes to its infected bacteria [57], was found in only 5 face samples, and in 23 of the gut samples at various levels of abundance (max. coverage= 17.9%, max. mapping reads= 372). We also found the *Shigella* phage SfIV, which aids the virulence of *Shigella flexneri* [58], in 25 of the gut samples and 8 of the face samples. Potentially pathogenic plasmids present only in the face microbiome are from opportunistic pathogens such as *Acinetobacter baumannii*, *Staphylococcus epidermidis*, which is usually part of the normal skin flora [59]. For example, the *Ochrobactrum anthropi* ATCC 49188 plasmid pOANT04 is present in 63 of the samples. *O. anthropi* is being increasingly recognized as a potentially problematic opportunistic and nosocomial human pathogen [60]. Examining the pathogenicity level of the differentially abundant bacterial pathogens, we found that 75.2% of the assigned pathogens are level 2 in the face samples (79 are level 2 and 26 are level 1), while 95.8% of those more abundant in the gut are level 2 (46 level 2 and 2 level 1).

Using the taxonomic annotations from the assembled genes, we found that the only protein from the MGmapper strict core identified in the highest number of face samples (20) and gut samples (36) was an uncharacterized protein from *Chlamydophila psittaci*, an avian pathogen that causes avian chlamydiosis, and epizootic outbreaks in mammals [61]. Among the viruses from this functional strict core we found the avian endogenous retrovirus EAV-HP and the avian leucosis virus. We also identified in higher abundance in the gut dataset *Trichuris trichiura*, causative of trichuriasis in humans [62,63] (max. mapping reads in the face samples= 942, max. mapping reads in the gut samples= 565,870), and *Eimeria brunetti*, causative of haemorrhagic intestinal coccidiosis in poultry [64] (max. mapping reads in the face samples= 30, max. mapping reads in the gut samples= 1,706). More abundant in the face dataset we identified the fly *Lucila cuprina* (mean abundance in the gut samples= 11,910, mean abundance in the face samples= 49,210), which causes sheep strike [65].

In order to highlight the importance of the cleaning service that vultures do to the environment, we searched for bacteria capable of zoonosis and sporulation [3] significantly more abundant in the gut microbiome compared to that of the face, since those are the ones that could potentially end back into the environment through the vulture’s faeces. We identified only 49 bacteria in the gut dataset with potential for sporulation, including *Fusobacterium necrophorum*, *Campylobacter jejuni*, and *Listeria monocytogenes* (Figure S9). Regarding the bacteria reported as zoonotic pathogens, we could only identify *Streptococcus suis*, a pig pathogen capable of transmission to humans from pigs [66]. Other identified bacteria with reported zoonotic capacity had very low abundance and were present in only one or two samples, so that they likely represent non-viable bacteria already dealt with by the vulture. **Stomach acidity protective evaluation**

To investigate the impact of the stomach acidity on the gut microbiota, and evaluate its acidity as a protection mechanism by the vulture physiology, we considered previously published measurements of pH from the stomachs with food contents from 13 wild black vultures [67]. Pure stomach acidity was 1.5-2.0 pH. A large standard deviation was observed in the stomach pH readings of the tested black vultures (2.0-5.6, x =3.8+1.25). In contrast, pH readings from the upper (5.6-7.3, x= 6.1+0.48) and lower intestine (5.9-7.0, x= 6.0+0.3) were more uniform, since the media itself is homogenized and has a high liquid content. To further test the hypothesis of a stomach acidity protection, we analysed the damage patterns and length distribution of the DNA reads mapped against the gut nr gene set catalogue. We did not identify any damage pattern in the DNA from the face nor the gut datasets, and did not identify a bimodal read length distribution in the gut dataset. Both face and gut DNA read length distributions were unimodal and similar (Figure S10). Interestingly, we identified the invertebrate *Clonorchis sinensis* in 22 of the face samples and 39 of the gut samples (66.6% of the facial and 83% of the gut samples). This liver fluke feeds on bile and causes problems in fat digestion, and it is able to reach the gut of the hosts given its acidity resistance [68].

## Discussion

### Microbiome composition and variability

The comparison of our metagenomic bacterial identifications to those by Roggenbuck *et al.* [21] confirm the consistency of the taxonomic identifications. Given that we aim at characterizing the vulture’s scavenging-related microbiome, in light of previous observations that the skin and gut microbiota of turkey and black vultures largely overlap [21], we combined the datasets of both vulture species into one. The results from comparing the number of identified taxa in each species prove that the microbiomes of these two vulture species are not statistically different (Figure S11), and validate their joint use. We identified a strikingly large taxonomic and functional variation in both gut and facial datasets (Figs. 3, 4, 6A). Examination of the PCA from the functional potential characterization suggests that the functional profile of the face and gut microbiomes are very similar, despite large intra-sample type variation. The suggestion is supported by the PCA of a down sampled dataset to the minimum of the distribution of each pathway class (Fig. 5), which further suggests that the relative abundance of the proteins rather than the presence/absence of them is one of the main factors distinguishing the face from the gut microbiome functional profile. In comparison to the face microbiome, the gut has less functional intra-sample variation (Table 1, *P*= 0.002). This is consistent with the fact that the face is the vulture’s first body part to make contact with the carcass. In general, the identified taxa from the gut and facial microbiomes can be classified as derived from *i*) host, such as *Methanobrevibacter smithii* in the gut, and *ii*) environment and carcass, e.g. *Xanthomonas* and *Actinobacillus pleuropneumoniae*. Given that the most abundant facial microbes can be associated to a variety of microbial attributes ranging from producers of antifungals to usual intestinal bacteria and plant and soil related bacteria, it is clear that there is a large environmental and carcass flora input to the vulture’s highly variable face microbiome. On the other hand, those most abundant in the gut are related mainly to intestinal or fecal bacteria, reflecting the digestive and more specialized functions expected to occur in the gut. Our data confirms previous PCR-based results [21] that identify *Clostridia* and *Fusobacteria* as dominant taxa in the gut microbiome (Figs. 2, 6C). As expected, there are clear traits in the gut microbiome for carrying out digestive and nutritional activities, which we identified in our datasets.

**Fig. 6.**
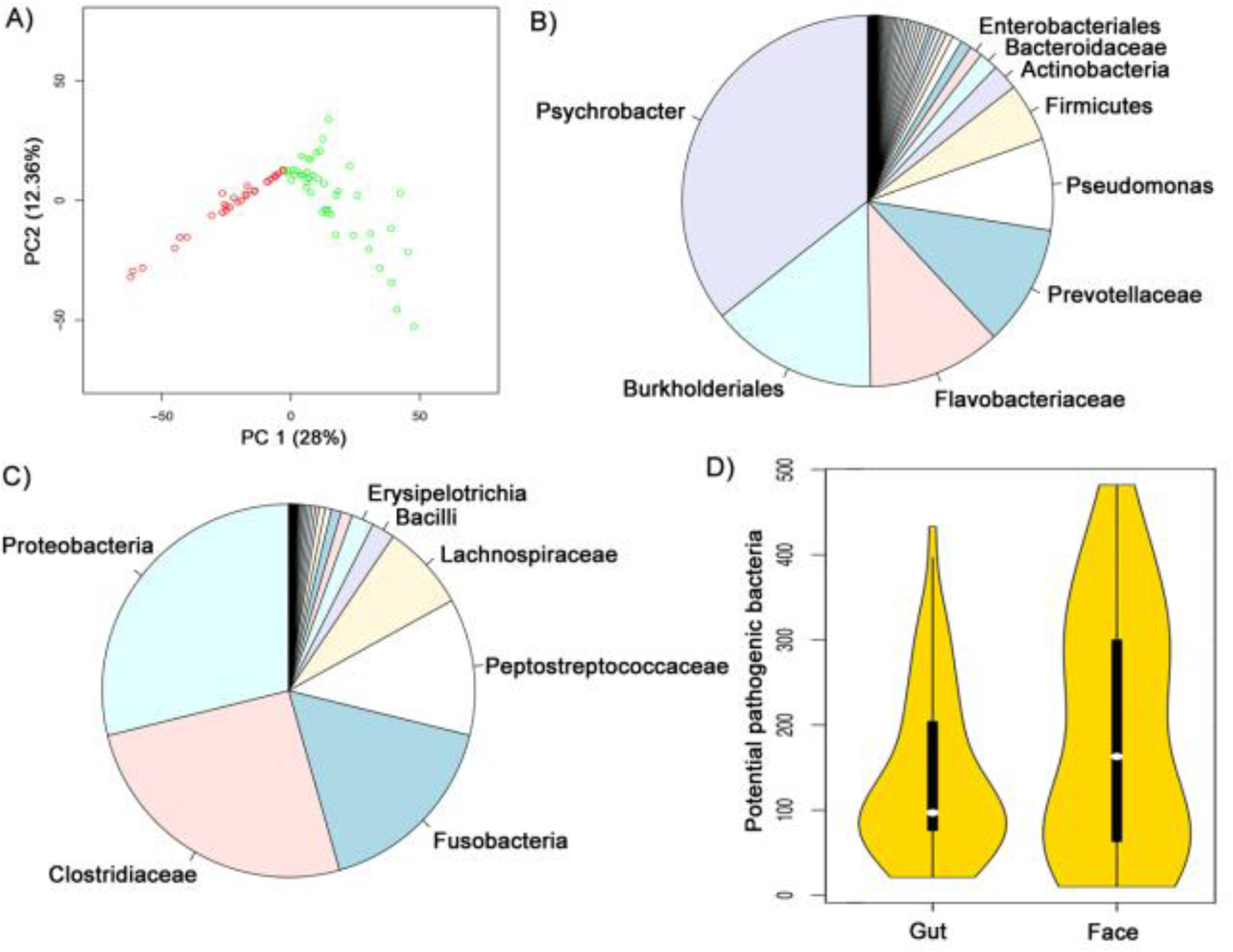
Vulture facial and gut microbiome composition. (**A**) Principal components (PCs) 1 (28% of the variation) and 2 (12.36%) of the abundance of all genes from all the KEGG metabolic classes together of the face (red) and gut (green) samples. (**B**) Taxonomic profile of the face and (**C**) gut microbiota. (**D**) Distribution of the number of identified potentially pathogenic bacteria in the gut and face datasets.

### Reduced core host microbiome

In order to differentiate the constant host microbiome from that derived from variable and external influences (i.e. microbes derived from the carcass and the environment), we defined two types of microbiome cores. A relaxed core containing those elements present in at least 50% of the samples, and a strict core with those present in at least 80% of the samples. We found that the relaxed core contains ~43% and ~22.7% of the facial and gut taxa, respectively, and the strict core only contains ~1% of the taxa in both gut and facial datasets (Supplementary Notes 1, 2). Notably, the distinction between carcass and established constant host-derived microbiome is complicated, even after the cores were defined. For example, the foodborne pathogen *Salmonella enterica* is present in the gut core (Fig. 6B). We additionally identified genes in the strict facial and gut microbiome functional cores that are related to putrescine, one of the main molecules produced in a carcass (Supplementary Notes 1, 3, Additional File 5). We found ~26x more habitat-specialized microbes in the face than in the gut microbiome (face= 8,373, gut= 320), most likely due to the fact that a mammalian corpse is a disturbance habitat that selects for a specialized microbial community [7]. Some of these community species likely derive from the carrion microflora, for example, we identified phenol degrading bacteria in the vulture facial skin, such as *Acinetobacter calcoaceticus* [69] (max. mapping reads in face samples= 478, max. mapping reads in gut samples= 6), for which we identified its gene coding for phenol 2-monooxygenase in the nr gene catalogue. Phenolic compounds can act against foodborne pathogens and spoilage bacteria [70], suggesting that they derive from carrion dwellers, adapted to their competitive environment, instead of being part of the core vulture facial skin microbiome. Thus, we suggest that the vulture microbiome is a result of its scavenging diet, with part of the carcass flora leaving a profound footprint in the vulture microbiome.

### Pathogenicity challenges

Given the identification of a strong carcass microbiota signature in the vulture microbiome, we next characterized all potential pathogens dealt with by the vultures. We define potential pathogens as taxa and functions that, while present in the vulture microbiome without conferring an apparently negative health effect, could be deadly for non-scavengers. Most of the significantly more abundant potential pathogens found in the facial microbiome (*P*< 0.05) are known to produce anthrax-like illnesses, periodontitis, pneumonia, and tuberculosis in mammals, while those found more abundantly in the gut are known to cause gastroenteritis, gas gangrene, food poisoning, and dysentery in humans (Figs. 6D, 7AB). We identified several pathogenic plasmids in the gut microbiome, such as the Shiga toxin 1-converting phage BP-4795, which transmits virulence genes to the infected *Escherichia coli* [57], as well as genes in the facial microbiome related to pathogenicity, such as hemolysins (Additional File 6). Our untargeted metagenomics approach also identified non-bacterial potential pathogens in the gut, such as the round worm *Trichuris trichiura*, causing trichuriasis in humans [63], and the apicomplexan parasite *Eimeria brunetti*, responsible for hemorrhagic intestinal coccidiosis in poultry [64]. These results highlight the health-challenging environment dealt by the vulture due to its scavenging diet.

### Stomach acidity protection

Most of the identified potential pathogens are restricted to few samples, and the abundance of the pathogens is not consistent across samples after counts normalization (Figure S12). These observations could be due to variations in the carcass flora, or efficient elimination of the potential pathogens by the vulture. Due to the previously reported very acid pH of some vulture species’ stomachs [3], the vulture stomach acidity has been suggested to serve as a filter of potential pathogens [8,21]. The stomach with food contents of the black vulture has a pH of 1.5-2.0 [67], a level common in mammals. Instrument readings of the pH are usually higher in a stomach with food contents since the gastric acid is sparse and diluted by the water content in the lumpy food items. Sequential and independent probe values in the stomach can often provide different readings, even when the probe is used in the same location. This is reflected in the large standard deviations observed in the previously reported pH readings of the tested black vultures [67]. The intestinal measurements were less acidic, and neutral in some occasions. Given that the carcass flora enters the vulture’s body mainly along with the ingested food items, the pH measurements suggest that the gastrointestinal acidity is not an efficient filter against all the potential pathogens present in a scavenging diet, rather it plays the general role of primary selection, which is not enough for all the potential pathogens in the carcass. In the comparison of the facial and gut microbiomes, we found that the face has more different species of potential pathogens than the intestine (*P*= 0.036) (Fig. 6D, Table 2), However, there is no statistical difference in their abundance (*P*= 0.82), and the intestine still harbours various potential pathogens. Furthermore, the lack of identification of a clear damage pattern in the reads could suggest that filtering of microbes occurs in the gastric chamber, thus the gut metagenome does not consist of both dead and living bacteria if the microbial filtering was to primarily occur there. This could also suggest that the gut taxa are able to withstand the physiological vulture’s gastrointestinal conditions, as supported by the identification of *Clonorchis sinensis.* Alternatively, the lack of DNA damage pattern and lack of length distribution bimodality could suggest that the identified predatory eukaryote *Adineta vaga* cleans the gut microbiome of dead bacteria and protozoa.

**Table 2.** Top 10 potential diseases causing bacteria identified in the face and gut.

| Pathogen | Disease | Facial mean abundance | Facial rank | Intestinal mean abundance | Intestinal rank |
| --- | --- | --- | --- | --- | --- |
| *Stenotrophomonas maltophilia* K279a | Bacteremia, bronchitis, pneumonia, urinary tract infection | 75,291.97 | 1 | 18,298 | 7 |
| *Clostridium perfringens* ATCC 13124 | Gas gangrene | 61,879.09 | 2 | 166,694.6 | 1 |
| *Plesiomonas shigelloides* 302-73 | Gastroenteritis | 48,642.03 | 3 | 134,119.1 | 2 |
| *Clostridium perfringens* str. 13 | Gas gangrene | 41,308.03 | 4 | 125,717.5 | 3 |
| *Pseudomonas fluorescens* A506 | Commensal (plant) | 31,324.91 | 5 | 271.1818 | 49 |
| *Clostridium perfringens* SM101 | Gas gangrene | 30,361.24 | 6 | 109,973.2 | 4 |
| *Acinetobacter lwoffii* SH145 | Nosocomial infections | 19,950.21 | 7 | 3.25 | 149 |
| *Aeromonas salmonicida* subsp. *salmonicida* A449 | Furunculosis | 14,347.45 | 8 | 45.45455 | 92 |
| *Acidovorax avenae* subsp. *avenae* ATCC 19860 | Bacterial leaf blight, brown stripe, red stripe | 13,111.21 | 9 | 998.75 | 30 |
| *Propionibacterium propionicum* F0230a | Commensal | 11,896.82 | 10 | 73.63636 | 80 |
| *Campylobacter lari* RM2100 | Gastroenteritis, diarrhea | 4,791.03 | 26 | 53,435.05 | 5 |
| *Campylobacter jejuni* subsp. *doylei* 269.97 | Bacteremia | 477.21 | 108 | 24,733.95 | 6 |
| *Campylobacter jejuni* RM1221 | Food poisoning | 327.45 | 120 | 13,136.93 | 9 |
| *Fusobacterium nucleatum* subsp. *nucleatum* ATCC 25586 | Periodontitis | 2166.03 | 57 | 13,456.57 | 8 |
| *Clostridium perfringens* E str. JGS1987 | Gastroenteritis, gas gangrene | 3,398.24 | 39 | 12,724.66 | 10 |

### Microbiome mediated protection

It has been shown that microbes provide protection to the host against pathogenic bacteria, thus we hypothesized that the vulture microbiome plays a protective role in terms of combating, preventing or maintaining the abundance in balance of potential pathogens. Accordingly, we identified functional and taxonomic protective elements that can be classified as related to *i*) beneficial bacterial taxa and functions, *ii*) phages, *iii*) predatory eukaryotes, and *iv*) colonization resistance (Fig. 7C, Supplementary Note 4).

**Fig. 7.**
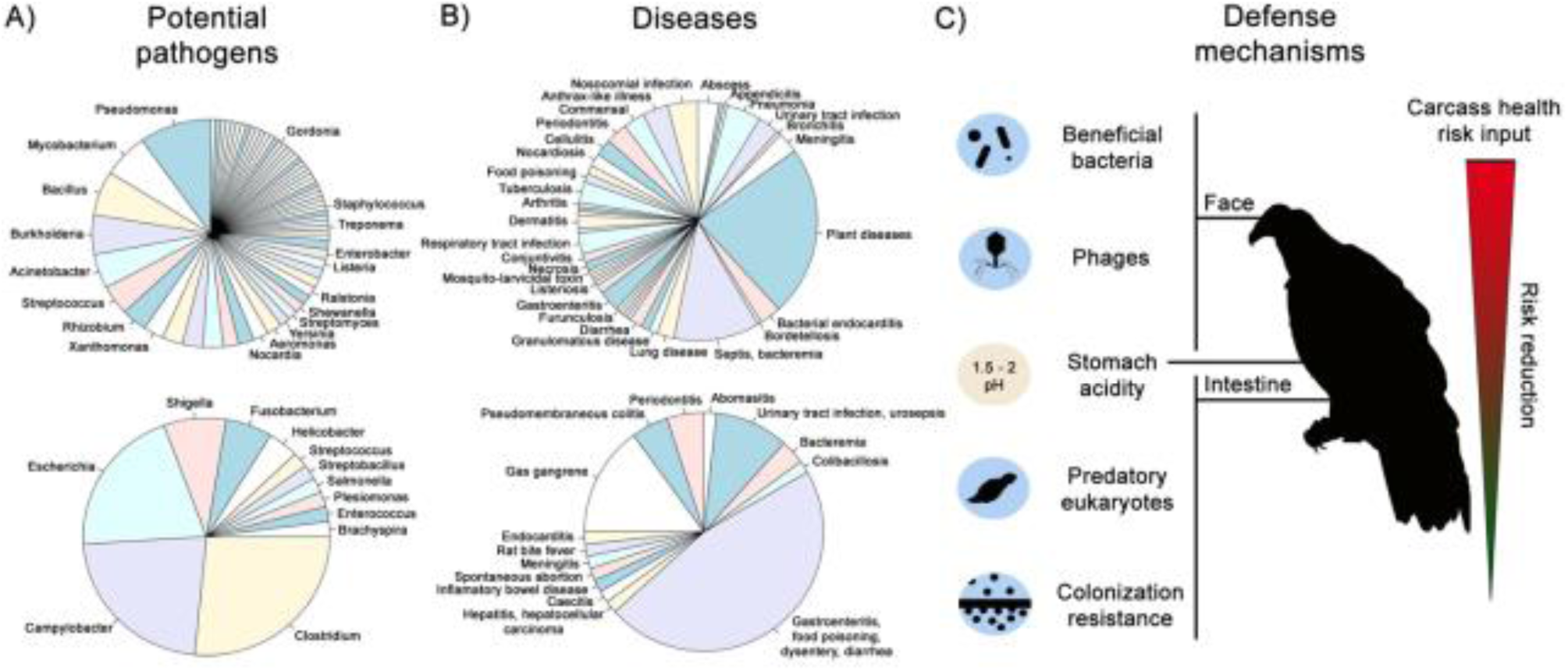
Health challenges faced by the vulture. **(A)** Vultures are confronted with a wide variety of carcass-derived microbes. (**B**) that pose serious pathogenic risks to non-scavenging species. (**C**) Different potential microbiome-mediated defence mechanisms account for the vulture’s ability to tolerate and reduce the health-risk potential that a carcass represents as it passes through its digestive system.

#### Beneficial bacterial taxa

Consistent with our microbiome-mediated protection hypothesis, we identified *Hylemonella gracilis* as part of the facial core, which has shown to prevent long term colonization by *Yersinia pestis* [71]. Other beneficial bacteria present in both gut and facial microbiomes include *Lactobacillus sakei*, an antilisterial bacteria [72]. We also identified several genes for the biosynthesis of antibiotics such as carbapenem, tetracycline, macrolides, and ansamycins, as well as resistance genes towards them (Additional File 6). The identification of insecticide, fungicide, and antiparasite related taxa and genes in the facial microbiome suggests protective mechanisms against possible eukaryotic pathogens present in the carcass (Additional File 6), such as *Pseudomonas entomophila* causing lethality in flies [73], for which we observed an insecticidal toxin SepC/Tcc class coding gene in the facial non-redundant gene set catalogue (Supplementary Note 5). The production of antibiotics to outcompete for resources is known in soil microbiomes, and recently similar strategies have been reported in the human nasal microbiome from commensal bacteria against pathogens [74]. Our results suggest that the vulture’s facial microbiome plays a similar defensive role. Regarding the gut microbiome, commensal *Clostridia* are known to play an important role in the production of butyrate that the colonocytes use [75]. Notably, *C. butyricum* is among the most abundant *Clostridia* in the gut microbiome.

#### Beneficial bacterial functions

Besides containing potential pathogenic microbes, carcasses also contain toxic and carcinogenic compounds [76], which pose health risks to the vulture, particularly to its bare skin face, which is in direct contact with such compounds. Among the bacteria identified in higher abundance in the facial microbiome is *Arthrobacter phenanthrenivorans*, which is able to degrade phenanthrene, a skin-irritating polycyclic aromatic hydrocarbon (PAH). PAH are xenobiotic pollutants with negative health-effects found to be emitted from animal carcass [77], and previously reported in high concentrations in other vulture species [78]. Interestingly, the largest intra-samples variation on the metabolism of xenobiotics biodegradation was in the facial dataset (Additional File 5), with PAHs degradation metabolism being the most abundant subclass from the xenobiotics degradation pathway in both face and gut. These findings suggest a microbiome protective role for the vulture against such compounds. In regards to the gut microbiome, the second most abundant *Fusobacteria* in the gut from the non-redundant gene set is the gut butyrate-producing *F. varium* [79], for which we also identified its gene formate C-acetyltransferase, which is involved in butanoate metabolism, as the most abundant gene in the gut dataset. Interestingly, besides the use of butyrate for the colonocytes, it has been shown that butyrate glycerides have antimicrobial activity against *C. perfringens* and *Salmonella typhimurium* [80].

#### Phage controlled pathogen abundance

Phages in the human gut microbiome have been shown to play a protective role and the increasing identification of antibiotic resistance genes in pathogenic bacteria has led to the proposition of using phages as alternative therapies [81]. Given the identification of potential antibiotic resistance genes in the vulture facial and gut microbiomes (Supplementary Note 6, Figures S13, S14), we investigated the possible role of phages in eliminating or balancing the abundance of potential pathogens. In the facial microbiome, *Clostridium* phages positively correlate with *Clostridium perfringes* and *Clostridium botulinum*, whereas in the gut we instead observed enterobacteria phages correlating to *Escherichia fergusonii* (Additional File 7). Furthermore, in the taxonomic annotations of the gut functional core, we identified the *Salmonella* phage SPN3US (Additional File 8), which has shown effective inhibition of *Salmonella enterica* [82]. From the facial functional strict core, the most abundant virus is phage BPP-1 (Additional File 8), which infects pathogenic *Bordetella* bacteria [83]. These findings show that the phage sets in both facial and gut microbiomes are related to the presence of the potential pathogens most abundant in the corresponding sample type and suggest that phages could represent, like phage therapy, an alternative defense mechanism for the control, and possibly elimination of potential pathogens [84] (Supplementary Note 5, Figure S15).

#### Predatory defense mechanism

In spite of being important elements in the gut microbiome, gut microbial eukaryotes remain largely unexplored. Thus, we investigated whether vulture gut microbial eukaryotes played any protective role. We identified the invertebrate *Adineta vaga*, which feeds on dead bacteria and protozoans, to be ~6.8 times more abundant in the gut core than in the facial core. This identification suggests that a predatory mechanism may be exploited for defense in the vulture’s gut.

#### Biofilm formation and colonization resistance

Biofilms are assemblages of microbes associated within a matrix composed of extracellular polymeric substances that facilitates their adhesion to the surface, protection against antimicrobials, and better nutrient acquisition. The abundance of *Fusobacteria* in the gut has been suggested to play a particularly relevant role in lumen biofilm formation in the gastrointestinal tract [85]. To explore this hypothesis, we searched for proteins related to biofilm formation (Additional File 8). In the core of the gut functional potential we identified biofilm related proteins from *Fusobacterium mortiferum*, such as rubrerythrin, as well as from *Clostridium perfringens*, such as UDP-glucuronic acidepimerase. Given that bacteria form biofilms in which they can thrive under different patterns of gene expression [86], this suggests that the identified potential pathogenic *Clostridia* and *Fusobacteria* from the gut microbiome might not pose pathogenic risks and instead offer colonization resistance against other external pathogens (Supplementary Note 5). To further explore the colonization resistance role of *Clostridia* and *Fusobacteria*, we examined the gut functional core for toxins with potential effects on the vulture. We identified only few potentially pathogenic toxin coding genes from *Fusobacterium* (Additional File 9), and *F. varium* has been shown to affect its human host in a beneficial manner by antagonizing colonization by pathogenic agents [87]. This suggests that an important role of the gut *Fusobacteria* could be the formation of biofilms and colonization resistance, without representing a serious pathogenic threat. We identified pathogenicity genes perfringolysin O and phospholipase C in the gut microbiome from *C. perfringens*. However, we also identified short chain fatty acids biosynthesis genes from *Clostridia*, which can provide protection against inflammatory responses [88]. Thus, we could classify the observed *Clostridia* into two types, *i*) the potentially pathogenic, mainly represented by *C. perfringens* and *ii*) the non-pathogenic, which may contribute to biofilm formation and health defense (Additional File 10).

## Conclusions

Our findings strongly suggest that these two species of vultures have adapted to their scavenging diet with the help of their facial and gut microbiomes. Surprisingly, most of their microbiome consists of a large variable pool of environmental and carcass-derived flora, with only a small set of constant inhabitants. In particular, we highlight the identification of defense mechanisms that are alternative to the use of antimicrobials, such as the use of predatory microbes, and the ironic protective nature of colonization resistance through biofilm formation by bacteria usually recognized as pathogenic to most mammals. The establishment of these mechanisms in the vulture microbiome highlights the important role vultures play in their ecosystem – performing an essential but underrated service by cleaning up carcasses that otherwise would spread pathogenic elements. In conclusion, our results highlight the importance of complementing genomic analyses with metagenomics on the host microbiome in order to obtain a clearer understanding of the host-microbial alliance that aids the evolution of extreme dietary adaptations.

## Declarations

### Ethics approval and consent to participate

We followed ethical guidelines and had ethics approval for the handling of our samples. Vulture trapping and euthanization procedures were approved by APHIS (Animal and Plant Health Inspection Service, USDA).

### Consent for publication

Not applicable.

### Availability of data and materials

The data reported in this paper are tabulated in the Supplementary Materials and the sequencing data will be archived in the NCBI SRA database.

### Authors’ contributions

MLZM performed the metagenomics analyses. KMV and MLZM performed the search for the resistance genes and potentially pathogenic microbes. GRG and MR provided the samples. MLZM, LHH, MTPG, GRG, MR, SR, and TSP interpreted the results. MLZM wrote the manuscript. LHH, MTPG, GRG, MR, SR, and TSP provided significant input on the manuscript drafting.

## Acknowledgements

We thank the Danish National High-Throughput DNA Sequencing Centre for the generation of the sequencing data. We are also grateful to George Pacheco and Lillian Anne Petersen for DNA extractions and library preparations. We also gratefully acknowledge the Danish National Supercomputer for Life Sciences – Computerome (computerome.dtu.dk) for the computational resources to perform the sequence analyses.

## References

1. Koenig R. Ornithology. Vulture research soars as the scavengers’ numbers decline. Science. 2006 Jun 16;312(5780):1591–2.

2. Margalida A, Colomer MÀ. Modelling the effects of sanitary policies on European vulture conservation. Sci Rep. 2012 Jan;2:753.

3. Houston DC, Cooper JE. The digestive tract of the whiteback griffon vulture and its role in disease transmission among wild ungulates. J Wildl Dis. 1975 Jul;11(3):306–13.

4. Anzen DH. Herbivores and the Number of Tree Species in Tropical Forests. Am Nat. 1970;104.940:501–28.

5. Vollaard EJ, Clasener HA. Colonization resistance. Antimicrob Agents Chemother. 1994 Mar;38(3):409–14.

6. Loeffler AG, Hart MN. Chapter 26 - Infectious Diseases. In: Publishers J& B, editor. Introduction to human disease: Pathophysiology for health professionals. 6th Editio. Burlington, MA 01803; 2014. p. 396–7.

7. Metcalf JL, Xu ZZ, Weiss S, Lax S, Van Treuren W, Hyde ER, et al. Microbial community assembly and metabolic function during mammalian corpse decomposition. Science (80-). 2015 Dec 10;351(6269):158–62.

8. Chung O, Jin S, Cho YS, Lim J, Kim H, Jho S, et al. The first whole genome and transcriptome of the cinereous vulture reveals adaptation in the gastric and immune defense systems and possible convergent evolution between the Old and New World vultures. Genome Biol. 2015 Oct 21;16(1):215.

9. Mateos-Hernndez L, Crespo E, la Fuente J de, de la Lastr JMP. Identification of Key Molecules Involved in the Protection of Vultures Against Pathogens and Toxins. In: An Integrated View of the Molecular Recognition and Toxinology - From Analytical Procedures to Biomedical Applications. InTech; 2013.

10. Ley RE, Lozupone CA, Hamady M, Knight R, Gordon JI. Worlds within worlds: evolution of the vertebrate gut microbiota. Nat Rev Microbiol. 2008 Oct;6(10):776–88.

11. Ley RE, Hamady M, Lozupone C, Turnbaugh PJ, Ramey RR, Bircher JS, et al. Evolution of mammals and their gut microbes. Science. 2008 Jun 20;320(5883):1647–51.

12. Semova I, Carten JD, Stombaugh J, Mackey LC, Knight R, Farber SA, et al. Microbiota regulate intestinal absorption and metabolism of fatty acids in the zebrafish. Cell Host Microbe. 2012 Sep 13;12(3):277–88.

13. Hajela N, Ramakrishna BS, Nair GB, Abraham P, Gopalan S, Ganguly NK. Gut microbiome, gut function, and probiotics: Implications for health. Indian J Gastroenterol. 2015 Mar;34(2):93–107.

14. O’Hara AM, Shanahan F. The gut flora as a forgotten organ. EMBO Rep. 2006 Jul;7(7):688–93.

15. Cryan JF, Dinan TG. Mind-altering microorganisms: the impact of the gut microbiota on brain and behaviour. Nat Rev Neurosci. Nature Publishing Group, a division of Macmillan Publishers Limited. All Rights Reserved.; 2012 Oct;13(10):701–12.

16. Hooper L V, Littman DR, Macpherson AJ. Interactions between the microbiota and the immune system. Science. 2012 Jun 8;336(6086):1268–73.

17. Buffie CG, Pamer EG. Microbiota-mediated colonization resistance against intestinal pathogens. Nat Rev Immunol. Nature Publishing Group, a division of Macmillan Publishers Limited. All Rights Reserved.; 2013 Nov;13(11):790–801.

18. Qin J, Li Y, Cai Z, Li S, Zhu J, Zhang F, et al. A metagenome-wide association study of gut microbiota in type 2 diabetes. Nature. 2012 Oct 4;490(7418):55–60.

19. Ley RE, Turnbaugh PJ, Klein S, Gordon JI. Microbial ecology: human gut microbes associated with obesity. Nature. 2006 Dec 21;444(7122):1022–3.

20. Sokol H, Pigneur B, Watterlot L, Lakhdari O, Bermúdez-Humarán LG, Gratadoux J-J, et al. Faecalibacterium prausnitzii is an anti-inflammatory commensal bacterium identified by gut microbiota analysis of Crohn disease patients. Proc Natl Acad Sci U S A. 2008 Oct 28;105(43):16731–6.

21. Roggenbuck M, Bærholm Schnell I, Blom N, Bælum J, Bertelsen MF, Pontén TS, et al. The microbiome of New World vultures. Nat Commun. Nature Publishing Group; 2014 Nov 25;5:5498.

22. Bolger AM, Lohse M, Usadel B. Trimmomatic: a flexible trimmer for Illumina sequence data. Bioinformatics. 2014 Apr 28;30(15):2114–20.

23. Zhang G, Li B, Li C, Gilbert M, Jarvis E, Consortium TAG, et al. The avian phylogenomics project data. GigaSci Database. 2014;

24. Kultima JR, Sunagawa S, Li J, Chen W, Chen H, Mende DR, et al. MOCAT: A Metagenomics Assembly and Gene Prediction Toolkit. 2012;7(10):1–6.

25. Petersen TN, Lukjancenko O, Thomsen MCF, Maddalena Sperotto M, Lund O, Møller Aarestrup F, et al. MGmapper: Reference based mapping and taxonomy annotation of metagenomics sequence reads. An L, editor. PLoS One. Public Library of Science; 2017 May 3;12(5):e0176469.

26. Li H, Durbin R. Fast and accurate short read alignment with Burrows-Wheeler transform. Bioinformatics. 2009 Jul 15;25(14):1754–60.

27. Nielsen HB, Almeida M, Juncker AS, Rasmussen S, Li J, Sunagawa S, et al. Identification and assembly of genomes and genetic elements in complex metagenomic samples without using reference genomes. Nat Biotechnol. 2014 Jul 6;32(8).

28. Peterson J, Garges S, Giovanni M, McInnes P, Wang L, Schloss JA, et al. The NIH Human Microbiome Project. Genome Res. 2009 Dec;19(12):2317–23.

29. Zankari E, Hasman H, Cosentino S, Vestergaard M, Rasmussen S, Lund O, et al. Identification of acquired antimicrobial resistance genes. J Antimicrob Chemother. 2012 Nov;67(11):2640–4.

30. DeSantis TZ, Hugenholtz P, Larsen N, Rojas M, Brodie EL, Keller K, et al. Greengenes, a chimera-checked 16S rRNA gene database and workbench compatible with ARB. Appl Environ Microbiol. 2006 Jul;72(7):5069–72.

31. Quast C, Pruesse E, Yilmaz P, Gerken J, Schweer T, Yarza P, et al. The SILVA ribosomal RNA gene database project: improved data processing and web-based tools. Nucleic Acids Res. 2013 Jan 1;41(Database issue):D590–6.

32. R Core Team. R: A language and environment for statistical computing. R Found Stat Comput Viena, Austria. 2013;

33. Huson DH, Weber N. Microbial community analysis using MEGAN. Methods Enzymol. 2013 Jan;531:465–85.

34. Edgar RC. Search and clustering orders of magnitude faster than BLAST. Bioinformatics. 2010 Oct 1;26(19):2460–1.

35. Gillespie JJ, Wattam AR, Cammer SA, Gabbard JL, Shukla MP, Dalay O, et al. PATRIC: the Comprehensive Bacterial Bioinformatics Resource with a Focus on Human Pathogenic Species. Infect Immun. 2011 Sep 6;79(11):4286–98.

36. van Belkum A. Classification of bacterial pathogens. COGEM Res Rep. 2011;1–113.

37. Ho Sui SJ, Fedynak A, Hsiao WWL, Langille MGI, Brinkman FSL. The association of virulence factors with genomic islands. PLoS One. 2009 Jan;4(12):e8094.

38. Buchfink B, Xie C, Huson DH. Fast and sensitive protein alignment using DIAMOND. Nat Methods. Nature Publishing Group, a division of Macmillan Publishers Limited. All Rights Reserved.; 2014 Nov 17;12(1):59–60.

39. Peng Y, Leung HCM, Yiu SM, Chin FYL. IDBA-UD: a de novo assembler for single-cell and metagenomic sequencing data with highly uneven depth. Bioinformatics. 2012 Jun 1;28(11):1420–8.

40. Hyatt D, Chen G-L, Locascio PF, Land ML, Larimer FW, Hauser LJ. Prodigal: prokaryotic gene recognition and translation initiation site identification. BMC Bioinformatics. 2010 Jan;11:119.

41. The universal protein resource (UniProt). Nucleic Acids Res. 2008 Jan;36(Database issue):D190–5.

42. Kanehisa M, Goto S. KEGG: Kyoto encyclopedia of genes and genomes. Nucleic Acids Res. 2000 Jan 1;28(1):27–30.

43. Luo R, Liu B, Xie Y, Li Z, Huang W, Yuan J, et al. SOAPdenovo2: an empirically improved memory-efficient short-read de novo assembler. Gigascience. 2012 Jan;1(1):18.

44. Gibson MK, Forsberg KJ, Dantas G. Improved annotation of antibiotic resistance determinants reveals microbial resistomes cluster by ecology. ISME J. International Society for Microbial Ecology; 2015 Jan;9(1):207–16.

45. Eddy SR. Accelerated Profile HMM Searches. PLoS Comput Biol. Public Library of Science; 2011 Oct 20;7(10):e1002195.

46. Jónsson H, Ginolhac A, Schubert M, Johnson PLF, Orlando L. mapDamage2.0: fast approximate Bayesian estimates of ancient DNA damage parameters. Bioinformatics. 2013 Jul 1;29(13):1682–4.

47. Martínez I, Lattimer JM, Hubach KL, Case JA, Yang J, Weber CG, et al. Gut microbiome composition is linked to whole grain-induced immunological improvements. ISME J. International Society for Microbial Ecology; 2013 Feb;7(2):269–80.

48. Rey FE, Faith JJ, Bain J, Muehlbauer MJ, Stevens RD, Newgard CB, et al. Dissecting the in vivo metabolic potential of two human gut acetogens. J Biol Chem. 2010 Jul 16;285(29):22082–90.

49. Leisner JJ, Laursen BG, Prévost H, Drider D, Dalgaard P. Carnobacterium: positive and negative effects in the environment and in foods. FEMS Microbiol Rev. Wiley-Blackwell; 2007 Sep 1;31(5):592–613.

50. Suzuki M, Yamamoto T, Kawai Y, Inoue N, Yamazaki K. Mode of action of piscicocin CS526 produced by Carnobacterium piscicola CS526. J Appl Microbiol. 2005 May;98(5):1146–51.

51. Murray WD, Khan AW, van den Berg L. Clostridium saccharolyticum sp. nov., a Saccharolytic Species from Sewage Sludge. Int J Syst Bacteriol. Microbiology Society; 1982 Jan 1;32(1):132–5.

52. He YL, Ding YF, Long YQ. Two cellulolytic Clostridium species: Clostridium cellulosi sp. nov. and Clostridium cellulofermentans sp. nov. Int J Syst Bacteriol. 1991 Apr;41(2):306–9.

53. Chassard C, Delmas E, Lawson PA, Bernalier-Donadille A. Bacteroides xylanisolvens sp. nov., a xylan-degrading bacterium isolated from human faeces. Int J Syst Evol Microbiol. 2008 Apr;58(Pt 4):1008–13.

54. George HA, Johnson JL, Moore WE, Holdeman L V, Chen JS. Acetone, Isopropanol, and Butanol Production by Clostridium beijerinckii (syn. Clostridium butylicum) and Clostridium aurantibutyricum. Appl Environ Microbiol. 1983 Mar;45(3):1160–3.

55. Madihah MS, Ariff AB, Khalil MS, Suraini AA, Karim MI. Anaerobic fermentation of gelatinized sago starch-derived sugars to acetone-1-butanol-ethanol solvent by Clostridium acetobutylicum. Folia Microbiol (Praha). 2001 Jan;46(3):197–204.

56. Frodyma ME, Downs D. The panE gene, encoding ketopantoate reductase, maps at 10 minutes and is allelic to apbA in Salmonella typhimurium. J Bacteriol. 1998 Sep;180(17):4757–9.

57. Creuzburg K, Recktenwald J, Kuhle V, Herold S, Hensel M, Schmidt H. The Shiga toxin 1-884 converting bacteriophage BP-4795 encodes an NleA-like type III effector protein. J Bacteriol. 2005 Dec 15;187(24):8494–8.

58. Jakhetia R, Talukder KA, Verma NK. Isolation, characterization and comparative genomics of bacteriophage SfIV: a novel serotype converting phage from Shigella flexneri. BMC Genomics. BioMed Central; 2013 Jan 3;14(1):677.

59. Grice EA, Kong HH, Renaud G, Young AC, Bouffard GG, Blakesley RW, et al. A diversity profile of the human skin microbiota. Genome Res. 2008 Jul;18(7):1043–50.

60. Chain PSG, Lang DM, Comerci DJ, Malfatti SA, Vergez LM, Shin M, et al. Genome of Ochrobactrum anthropi ATCC 49188T, a Versatile Opportunistic Pathogen and Symbiont of Several Eukaryotic Hosts. J Bacteriol. 2011 Jun 17;193(16):4274–5.

61. Harkinezhad T, Geens T, Vanrompay D. Chlamydophila psittaci infections in birds: a review with emphasis on zoonotic consequences. Vet Microbiol. 2009 Mar 16;135(1–2):68–77.

62. Klementowicz JE, Travis MA, Grencis RK. Trichuris muris: a model of gastrointestinal parasite infection. Semin Immunopathol. 2012 Nov;34(6):815–28.

63. Jung RC, Beaver PC. Clinical observations on Trichocephalus trichiurus (whipworm) infestation in children. Pediatrics. 1951 Oct;8(4):548–57.

64. Shirley MW, Smith AL, Tomley FM. The biology of avian Eimeria with an emphasis on their control by vaccination. Adv Parasitol. 2005 Jan;60:285–330.

65. East IJ, Eisemann CH. Vaccination against Lucilia cuprina: the causative agent of sheep blowfly strike. Immunol Cell Biol. Australasian Society for Immunology Inc.; 1993 Oct;71 (Pt 5)(5):453–62.

66. Wertheim HFL, Nghia HDT, Taylor W, Schultsz C. Streptococcus suis: an emerging human pathogen. Clin Infect Dis. 2009 Mar 1;48(5):617–25.

67. Graves G. Field measurements of gastrointestinal pH of New World vultures in Guyana. J Raptor Res. 2016;Submitted.

68. Hou PC. The pathology of Clonorchis sinensis infestation of the liver. J Pathol Bacteriol. 1955 Jul;70(1):53–64.

69. Paller G, Hommel RK, Kleber HP. Phenol degradation by Acinetobacter calcoaceticus NCIB 8250. J Basic Microbiol. 1995 Jan;35(5):325–35.

70. Nazzaro F, Fratianni F, De Martino L, Coppola R, De Feo V. Effect of essential oils on pathogenic bacteria. Pharmaceuticals (Basel). 2013 Jan;6(12):1451–74.

71. Pawlowski DR, Raslawsky A, Siebert G, J Metzger D. Identification of Hylemonella gracilis as an Antagonist of Yersinia pestis Persistence. J Bioterror Biodef. OMICS International; 2011 Feb 10;2(S3).

72. Bredholt S, Nesbakken T, Holck A. Industrial application of an antilisterial strain of Lactobacillus sakei as a protective culture and its effect on the sensory acceptability of cooked, sliced, vacuum-packaged meats. Int J Food Microbiol. 2001 Jun 15;66(3):191–6.

73. Vodovar N, Vinals M, Liehl P, Basset A, Degrouard J, Spellman P, et al. Drosophila host defense after oral infection by an entomopathogenic Pseudomonas species. Proc Natl Acad Sci U S A. 2005 Aug 9;102(32):11414–9.

74. Zipperer A, Konnerth MC, Laux C, Berscheid A, Janek D, Weidenmaier C, et al. Human commensals producing a novel antibiotic impair pathogen colonization. Nature. Nature Research; 2016 Jul 27;535(7613):511–6.

75. Pryde SE, Duncan SH, Hold GL, Stewart CS, Flint HJ. The microbiology of butyrate formation in the human colon. FEMS Microbiol Lett. 2002 Dec 17;217(2):133–9.

76. Phan N-T, Kim K-H, Jeon E-C, Kim U-H, Sohn JR, Pandey SK. Analysis of volatile organic compounds released during food decaying processes. Environ Monit Assess. 2012 Mar;184(3):1683–92.

77. Chen S-J, Hsieh L-T, Chiu S-C. Emission of polycyclic aromatic hydrocarbons from animal carcass incinerators. Sci Total Environ. 2003;313(1):61–76.

78. Dhananjayan V, Muralidharan S. Levels of polycyclic aromatic hydrocarbons, polychlorinated biphenyls, and organochlorine pesticides in various tissues of white-backed vulture in India. Biomed Res Int. 2013 Jan;2013:190353.

79. Dzink JL, Socransky SS. Amino acid utilization by Fusobacterium nucleatum grown in a chemically defined medium. Oral Microbiol Immunol. 1990 Jun;5(3):172–4.

80. Namkung H, Yu H, Gong J, Leeson S. Antimicrobial activity of butyrate glycerides toward Salmonella Typhimurium and Clostridium perfringens. Poult Sci. 2011 Oct;90(10):2217–22.

81. Sulakvelidze A, Alavidze Z, Morris JG. Bacteriophage therapy. Antimicrob Agents Chemother. American Society for Microbiology (ASM); 2001 Mar;45(3):649–59.

82. Kagawa H, Ono N, Enomoto M, Komeda Y. Bacteriophage chi sensitivity and motility of Escherichia coli K-12 and Salmonella typhimurium Fla-mutants possessing the hook structure. J Bacteriol. 1984 Feb;157(2):649–54.

83. Liu M, Deora R, Doulatov SR, Gingery M, Eiserling FA, Preston A, et al. Reverse transcriptase-mediated tropism switching in Bordetella bacteriophage. Science. American Association for the Advancement of Science; 2002 Mar 15;295(5562):2091–4.

84. Focà A, Liberto MC, Quirino A, Marascio N, Zicca E, Pavia G, et al. Gut Inflammation and Immunity: What Is the Role of the Human Gut Virome? Mediators Inflamm. Hindawi Publishing Corporation; 2015;2015:1–7.

85. Keenan SW, Engel AS, Elsey RM. The alligator gut microbiome and implications for archosaur symbioses. Sci Rep. Nature Publishing Group; 2013 Jan 7;3:2877.

86. Prigent-Combaret C, Vidal O, Dorel C, Lejeune P. Abiotic surface sensing and biofilm-dependent regulation of gene expression in Escherichia coli. J Bacteriol. 1999 Oct;181(19):5993–6002.

87. Ushijima T, Ozaki Y. Potent antagonism of Escherichia coli, Bacteroides ovatus, Fusobacterium varium, and Enterococcus faecalis, alone or in combination, for enteropathogens in anaerobic continuous flow cultures. J Med Microbiol. 1986 Sep;22(2):157–63.

88. Vinolo MAR, Rodrigues HG, Nachbar RT, Curi R. Regulation of inflammation by short chain fatty acids. Nutrients. Multidisciplinary Digital Publishing Institute (MDPI); 2011 Oct;3(10):858–76.

